# Enhanced microglial dynamics and paucity of tau seeding in the amyloid plaque microenvironment contributes to cognitive resilience in Alzheimer’s disease

**DOI:** 10.1101/2023.07.27.550884

**Authors:** Nur Jury-Garfe, Yanwen You, Pablo Martínez, Javier Redding-Ochoa, Hande Karahan, Travis S. Johnson, Jie Zhang, Jungsu Kim, Juan C. Troncoso, Cristian A. Lasagna-Reeves

## Abstract

Asymptomatic Alzheimer’s disease (AsymAD) describes the status of subjects with preserved cognition but with identifiable Alzheimer’s disease (AD) brain pathology (i.e. Aβ-amyloid deposits, neuritic plaques, and neurofibrillary tangles) at autopsy. In this study, we investigated the postmortem brains of a cohort of AsymAD cases to gain insight into the underlying mechanisms of resilience to AD pathology and cognitive decline. Our results showed that AsymAD cases exhibit an enrichment of core plaques and decreased filamentous plaque accumulation, as well as an increase in microglia surrounding this last type. In AsymAD cases we found less pathological tau aggregation in dystrophic neurites compared to AD and tau seeding activity comparable to healthy control subjects. We used spatial transcriptomics to further characterize the plaque niche and found autophagy, endocytosis, and phagocytosis within the top upregulated pathways in the AsymAD plaque niche, but not in AD. Furthermore, we found ARP2, an actin-based motility protein crucial to initiate the formation of new actin filaments, increased within microglia in the proximity of amyloid plaques in AsymAD. Our findings support that the amyloid-plaque microenvironment in AsymAD cases is characterized by microglia with highly efficient actin-based cell motility mechanisms and decreased tau seeding compared to AD. These two mechanisms can potentially provide protection against the toxic cascade initiated by Aβ that preserves brain health and slows down the progression of AD pathology.

## Introduction

Aβ-amyloid plaques and neurofibrillary tau tangles (NFTs) have been causally related to the cognitive manifestations of Alzheimer’s disease (AD)^1^ for decades. However, several studies have revealed the existence of aged individuals harboring a high burden of brain lesions at autopsy while remaining cognitively intact, indicating resilience to AD pathology^2–11^. These individuals have comparable neuritic plaque scores (CERAD)^12^ and Braak NFT stages^13^ to those of demented AD cases at autopsy, and the literature refers them as resilient^14^, non-demented individuals with AD pathology (NDAN)^15^ or asymptomatic AD (AsymAD)^10, 16^. We will use the last term in the present study. Several reports provide insight into the resistance of AsymAD subjects to cognitive decline. Specifically, studies have demonstrated that AsymAD brains exhibit no signs of notorious synaptic or neuron deterioration^16–18^, and intriguingly, even show larger nuclei and cellular sizes than age-matched controls^19^. Additionally and contrary to brains of AD demented patients, there is no evidence of phosphorylated tau accumulation within the synapses of AsymAD brains^15, 18^. On the other hand, AsymAD cases have been found to exhibit a distinct neuroinflammatory profile compared to AD brains, with decreased number of microglia and astrocytes^18^, as well as low levels of pro-inflammatory cytokines and increased anti-inflammatory cytokines^20^. Advancements in omics and large cohort data set analyses have also enabled the identification of potential cell signatures and molecular mechanisms of resilience, including high processing of energetic pathways involving mitochondrial metabolism and glycolysis, axonal and dendritic growth, and general increase of protein processing^21–23^.

Despite multiple studies using whole brain approaches, bulk proteomics and transcriptomics aimed at understanding how synaptic preservation and neuron survival are achieved in AsymAD brains, the molecular mechanisms underlying the resilience in the presence of NFTs and amyloid plaques are still not well understood. Growing evidence suggests that investigating the AD pathology with a spatial approach is important to understand molecular pathways involved in neurodegeneration. Moreover, the microenvironment in Aβ amyloid plaques play a crucial role in the Aβ-mediated neuroinflammation and tau pathogenesis in AD mice models^24–26^. These studies found that Aβ plaques create a unique environment that facilitates the rapid amplification of proteopathic tau seeds into large tau aggregates, initially appearing as dystrophic neurites surrounding Aβ plaques (NP-tau) followed by the formation and spread of tau aggregates^24^. Moreover, an efficient microglia clustering around Aβ plaques mitigates amyloid-driven tau seeding^27^. In the context of AD resilience, one study showed that the area surrounding NFTs in the hippocampus of AsymAD individuals exhibits lower levels of proteins associated with inflammation, oxidative stress, and high energy demands when compared to AD subjects^28^. Additionally, AsymAD cases show a significant upregulation of phagocytic microglia that helps to remove damaged synapses as a protective mechanism^29^. Taken together, these data point at the local milieu of Aβ amyloid as the crucial starting point of tau-driven synapse damage in human brains. Nevertheless, further studies are needed to provide a detailed insight into the mechanisms underlying Aβ plaque-associated microglial reactivity and tau pathogenesis in the context of resilience to AD pathology.

Herein, using postmortem brain samples from AsymAD, demented AD cases, and age-matched controls individuals, we performed a detailed histological and biochemical characterization of Aβ amyloid plaques and their cellular microenvironment, including microglia and astrocytes activation and tau pathology. We also performed spatial whole transcriptomics analyses to identify and characterize neuroprotective mechanisms operating in the amyloid plaque microenvironment and their potential contribution to cognitive resilience in AsymAD cases. We found that in AsymAD cases there is an enrichment of core-plaques in compared to AD. In contrast, filamentous plaques are predominant in AD. We also observed a strong engagement of microglia around filamentous plaques, with a concomitant strong reduction in NP-tau and tau-seeding activity in AsymAD in comparison to AD. Using spatial whole transcriptomics, we further demonstrated that in the amyloid-plaque microenvironment of AsymAD individuals, microglia have significant upregulation of actin-based motility genes. This upregulation may heighten microtubules dynamics, facilitating efficient migration towards the vicinity of the plaque and promoting elongation of microglial branches to enhance its engagement with the plaque. Furthermore, once microglia in AsymAD brains embrace the amyloid plaque, they may have more efficient autophagy mechanisms to degrade amyloid in comparison with AD cases. Understanding the local drivers of resilience to Alzheimer’s pathology may provide valuable insights into developing interventions to halt neuronal and synaptic damage and prevent the clinical manifestations of AD.

## Materials and methods

### Subjects and clinical-neuropathological classification

We examined middle frontal gyrus (MFG) tissue sections from individuals with histopathologic findings of Alzheimer’s disease (i.e., amyloid plaques and NFTs) and healthy aged-matched controls (Table 1). The cognitive status before death was obtained from detailed neuropsychological assessments and a diagnosis of dementia was defined according to the standard Mini-Mental State Examination (MMSE), Clinical Dementia Rating (CDR) scores and expert discussion at clinical conferences. The cognitive status and neuropathologic data were provided by the Johns Hopkins Brain Resource Center (BRC). Based on clinical and neuropathological data previously published^16, 19, 30, 31^ the brains were classified into aged-matched controls, asymptomatic for Alzheimer’s disease (AsymAD) and Alzheimer’s dementia (AD). The experimental groups have similar ages, male/female distribution, years of education, and number of APOE e4 alleles.

**TABLE 1:**
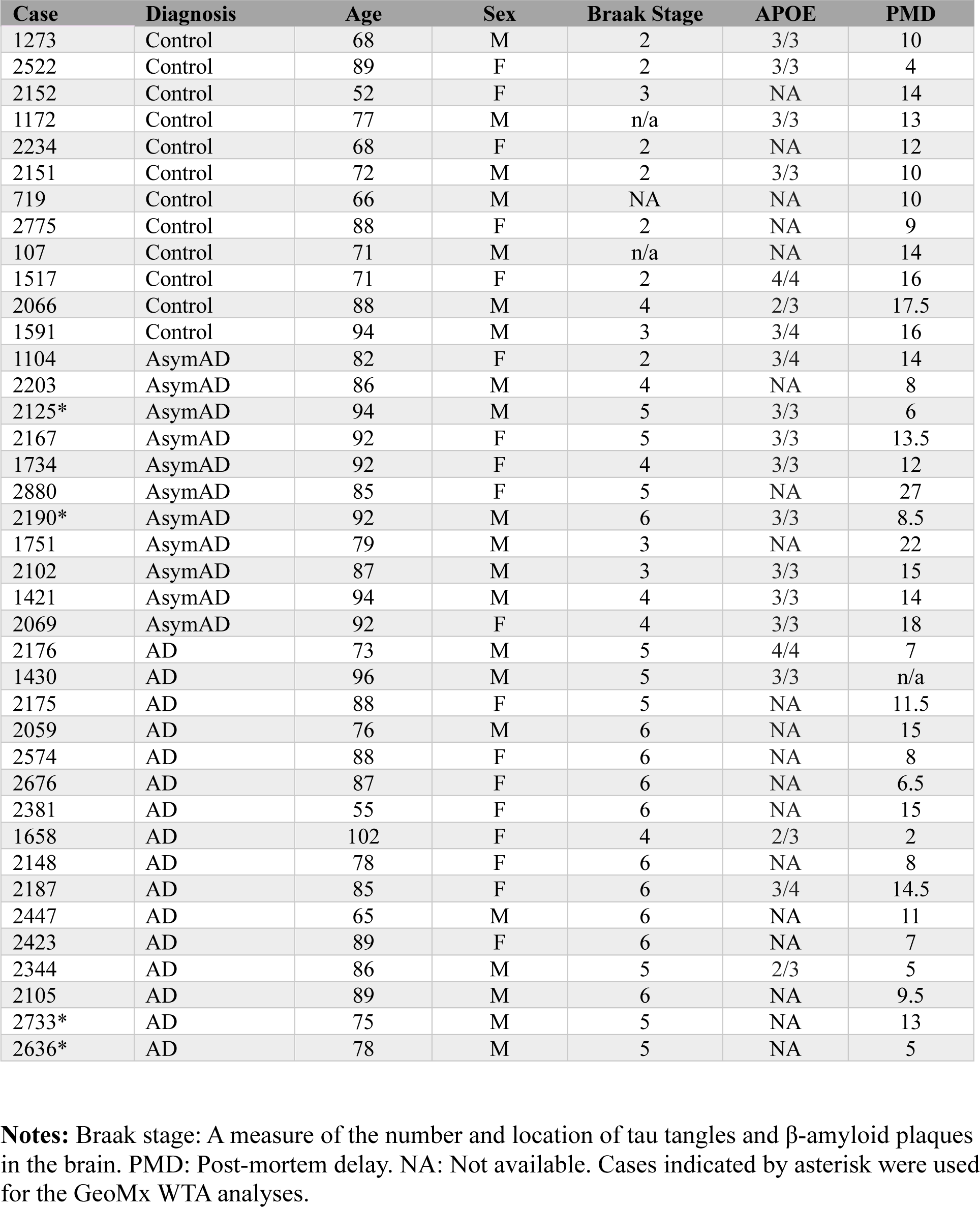
Clinical data.

### Tissue processing and neuropathologic evaluations

All brains were examined in the Division of Neuropathology at Johns Hopkins University. After weighing and external brain examination was performed, the brain is hemisected through the midline and the right cerebral hemisphere was cut serially in 1cm-thick coronal slabs. For diagnostic purposes, tissue blocks were fixed in 10% neutral buffered formalin, processed overnight, and embedded in paraffin. The tissue blocks were cut at 5 μm and stained with hematoxylin and eosin. Selected sections were stained with the Hirano silver method^32^ or treated with H_2_O_2_, and blocked with 3% normal goat serum in Tris-buffered saline for immunostaining with the phospho-tau (Ser202, Thr205) AT8 antibody (1:200, MN1020, Invitrogen).

### Immunofluorescence in post-mortem human tissue

Formalin-fixed, paraffin-embedded (FFPE) MFG 5 μm thick sections were deparaffinized in xylene, rehydrated in an ethanol gradient (100% to 30%) and washed with deionized water. Then, the sections were heated to 95 °C in 1X EDTA Buffer, pH 8.5 antigen retrieval solution (E1161, Sigma-Aldrich) for 20 min in a thermoregulated bath. For ARP2 and LAMP2, two additional steps of incubation with 50% Methanol during 15 min. and 1 min. of incubation with proteinase K (1 mg/ml) were performed. After washing twice with TBS of 5 min each, the sections were blocked with 3% goat serum/ 3% BSA in TBS 1x 0.01% Triton X-100 for 1 h at RT. Sections were then incubated overnight at 4 °C with the following primary antibodies: anti-IBA1 (1:300, 019-19741, Wako), anti-GFAP (1:300, ab1218, Abcam), AT8 (1:100, MN1020, Invitrogen), anti-ARP2 (E-12) (1:100, sc-166103, Santa Cruz) and anti-LAMP2 (H4B4) (1:300, ab25631, Abcam). The next day, sections were quickly washed three times in TBS and incubated for 2 h with a 1:500 ratio of Alexa Fluor antibodies; goat anti-rabbit Alexa Fluor 488 (A11008, Invitrogen), goat anti-mouse Alexa Fluor 488 (A32723, Invitrogen) diluted in blocking solution followed by three washes with TBS (5 min each). When necessary, amyloid structures were stained with 1% thioflavin S (diluted in 50% TBS/Ethanol) or NucBlue (1:100, R37605, Invitrogen) for 20 min at RT followed by five washes with TBS of 5 min each. Finally, sections were incubated with TrueBlack (23007, Biotium) for 1 min and then washed three times in TBS.

### Image Analysis

Brightfield images were acquired using the BH-2 Olympus microscope. Fluorescence imaging was performed using the Nikon A1-R laser scanning confocal microscope coupled with Nikon AR software v.5.21.03. Post processing and analysis was done using ImageJ (National Institutes of Health, v1.53c). Phospho-tau quantification in neurites using brightfield images was done as previously published^33^. For plaque classification analyses, 15–20 μm z-stacks were imaged at a 60X magnification and circularity analysis on thioflavin S plaques was performed using the ‘‘Shape Descriptors’’ plugin in ImageJ. Cutoffs for plaque circularity were defined as previously published^26, 34^, where filamentous plaques had a circularity score of 0.00–0.14 and compact plaques had circularities greater than 0.30. Plaques falling with the circularity scores range of 0.15–0.28 were classified as displaying intermediate phenotypes. The quantification of IBA1 and GFAP coverage around plaques was done as previously published^35^ with some modifications; regions-of-interest (ROIs) were traced along 50 μm of thioflavin S plaque perimeter in serial sections. Defined ROIs were applied to the IBA1/GFAP channels, and the percentage of immunoreactivity in the area within the ROI was quantified. Between 10-30 plaques in total were quantified. For NP-tau quantifications; AT8 positive puncta surrounded amyloid plaques (within a 50 μm perimeter) were quantified using the ‘‘analyzed particles’’ tool. To avoid comparisons between cases with extreme tau pathology, AD and AsymAD cases with a Braak and Braak scores of 4–5 were used to quantify NP-tau and AT8 neuritic density.

For ARP2 and LAMP2 quantifications, ROIs were traced along 50 μm of the plaque core, followed by the quantification of ARP2 immunoreactivity within the selected area. In addition, to quantify ARP2/LAMP2 exclusively in IBA1 positive staining, IBA1 mask selection was applied and ARP2/LAMP2 immunoreactivity and positive area were quantified. Colocalization was quantified by Pearson correlation coefficient using the JACoP plugin in ImageJ Fiji. Image fields more than 100 μm far from amyloid plaques were considered ‘‘plaque-free areas’’.

### Western blot analysis

Postmortem MFG tissues were homogenized in TBS buffer at a ratio of 1:10 (wt/vol) with Pierce Protease and Phosphatase Inhibitor Cocktail (A32965, ThermoScientific) on ice. Tissue lysate was sonicated and then centrifuged at maximum speed for 15 min at 4 °C. Protein concentrations were measured using the BCA protein assay kit (Bio-Rad Laboratories, Inc.). Electrophoresis was performed using 30 μg of protein lysates, resolved in a 4–12% SDS-PAGE gel (CriterionTM TGXTM, Bio-Rad Laboratories, Inc.) and transferred to a nitrocellulose membrane (Immobilon®-P, Millipore) that was blocked with 5% BSA in TBS with 0.01% tween, followed by overnight incubation of primary antibodies; HT7 (1:300, MN1000, Thermo Fisher), AT8 (1:1000, MN1020, Invitrogen) and PHF1 (1:1000, Peter Davies antibodies) diluted in the blocking solution. Horseradish peroxidase (HRP) secondary antibodies (goat anti-mouse HRP conjugated (1:10,000, 626820, Invitrogen) were incubated for 2 h at RT and the proteins were detected with Supersignal West Pico (34580, Thermo Scientific) and imaged by using iBright 1500 (Invitrogen). Western blots were analyzed using ImageJ Fiji.

### Meso Scale Discovery (MSD) of Aβ40 and Aβ42 levels

For Aβ40 and Aβ42 detection, V-PLEX Plus Aβ Peptide Panel 1 (6E10) Kit (K15200E; Meso Scale Discovery, MSD) was used. TBS-Soluble Aβ40 and Aβ42 levels were measured in MFG fractions of brain samples. The assay was performed according to the manufacturer’s instructions. Briefly, the plate was blocked with MSD Diluent 35 for 1h at room temperature (RT) with shaking at 700 rpm and washed three times with PBS-Tween (PBS-T). SULFO-TAG 6E10 detection antibody and samples or calibrators were loaded into the plate and incubated at RT for 2 h with shaking at 700 rpm. After three washing steps with PBS-T, MSD Read Buffer was added into the wells and the electrochemiluminescent signals were measured using a MESO QuickPlex SQ 120 Imager. The concentrations were normalized by total protein concentrations for each sample.

### Size exclusion chromatography (SEC)

SEC was performed as previously described^36^. Briefly, the column was equilibrated, and samples were clarified by centrifugation at 10,000 *g* for 10 min. Protein concentration from frozen samples was quantified by BCA assay, and 1–5 mg total protein of supernatant was taken for separation. The supernatant was concentrated with a 0.5 ml 3K Amicon centrifugal filter (UFC5003, Millipore Sigma) to ∼200 µl, then loaded onto the column via sample loop injection. Starting from injection, 1 ml fractions were collected into tubes containing EDTA-free protease inhibitor (11873580001, Roche) at a flow rate of 0.3 ml min^−1^.

### Human tau ELISA

ELISA was performed on total and SEC fractions using Tau (Total) Human ELISA Kit (KHB0041, Invitrogen) by following the directions provided by the manufacturer. Lysates were diluted 1:50,000 in blocking buffer. F7–F14 were diluted at a ratio of 1:2,000 in blocking buffer. F15– F22 were diluted at a ratio of 1:20,000 in blocking buffer.

### Tau-seeding assay

The seeding assay was performed as previously described^36, 37^. Briefly, TauRD P301S FRET Biosensor cells (CRL-3275, American Type Culture Collection (ATCC)) were plated at 35,000 cells per well in 130 µl medium in a 96-well plate, then incubated at 37 °C overnight. The next day, cells were transfected with total protein lysates and SEC fractions from control, AD and AsymAD cases (20 µg total protein per well). After harvest, flow cytometry was conducted with a BD LSR Fortessa X-20 with a High Throughput Sampler, using the BD FACS Diva v8.0 software. FlowJo v10.0 was used for data analysis. Seeding was quantified by integrated FRET density, defined as the product of the percentage of FRET-positive cells and median fluorescence intensity of FRET-positive cells.

### NanoString GeoMx^TM^ human whole transcriptome atlas (HuWTA)

Slide preparation was performed following the manufacturer’s instructions in the GeoMx WTA kit. Briefly, FFPE MFG sections were deparaffinized and rehydrated followed by antigen and target retrieval and *in situ* hybridization at 37°C during 16-24 h. Next day, slides were washed and incubated with the morphology markers; β-Amyloid (D54D2) Alexa Fluor 594 conjugated (1:100, Cell Signaling, 35363) and the nuclear marker Syto 13 (1:50, Nanostring, 121301310) during 1 h. Slides were loaded in the GeoMx Digital Spatial Profiler (DSP) instrument and scanned to capture fluorescent images used to select ROIs, a 50-μm in diameter circle was selected as the center ROI surrounding each plaque, a total of 16-18 ROIs per case (*n=2,* per condition) were selected within the gray matter. UV-cleaved oligonucleotides from each spatially resolved ROI were aspirated and collected in a 96-well collection plate to perform library prep with Seq Code primers and sequenced on an NextSeq500 sequencer instrument (Illumina). Digital count conversion files (DCC) were obtained using the Illumina DRAGEN Sever v4. DCC files were transfer to the GeoMx DSP Analysis suite v.3.0.0.109 and data quality control (QC) was performed followed by normalization. The analysis pipeline was done using the GeoMx DSP user manual (MAN-10154-01). Differential gene expression (DEGs) analyses (*p* < 0.05 with multiple testing correction, fold-change > 1.5) and pathways enrichment analyses using a total of 696 DEGs found in AsymAD cases, were performed.

### Statistical analyses

All statistical analysis and graph designs were performed using GraphPad Prism v9.5.0 (525). Data were first analyzed for normality (Shapiro-Wilk test) followed by statistical tests. Results in column graphs represent the mean ± S.E.M. For histology, immunofluorescence, and biochemical experiments, a Student’s *t*-test was performed to compare two groups, One-way ANOVA followed by multiple comparisons was employed for the comparisons of three groups and two-way ANOVA to analyze two variables simultaneously. For all tests, a *p-*value of 0.05 was used to determine statistical significance. When data were not normally distributed, Mann Whitney test was applied comparing two groups. For GeoMx HuWTA analyses, the *p* values were adjusted for multiple analyses using Benjamini-Hochberg procedure with a false discovery rate (FDR) of 0.01. In all quantifications, sex was considered as a biological variable. Data collection and analysis were performed blind to the conditions of the experiments.

## Results

### Differential plaque phenotypic distribution and Aβ42/40 ratio in AsymAD cases compared to Alzheimer’s disease cases

Given that autopsies of AsymAD and AD subjects reveal the presence of comparable amyloid plaques and neurofibrillary tangles (NFTs) **(Supplementary Fig. 1)**, we aimed to investigate if there were any differences in plaque morphology between these cases. To do so, we evaluated amyloid plaque phenotype in the MFG of thioflavin S stained sections **(Fig. 1A)**. Our analysis of the total number of amyloid plaques within the MFG did not reveal significant differences between AD and AsymAD cases **(Fig. 1B)**. Using previously described plaque classifications^26, 34^ and circularity analysis to distinguish compact plaques from plaques with a filamentous or an intermediate morphology **(Fig. 1C)**, we observed that AsymAD brains showed a significant reduction in the proportion of filamentous plaques, with a concomitant increase in compact plaques when compared to demented AD **(Fig. 1D)**. Interestingly, it has been previously reported that filamentous plaques are neurotoxic whereas compact dense-core are considered relatively benign^38–40^. We then evaluated soluble Aβ_42_ and Aβ_40_ peptides using MSD ELISA. Our results showed no differences in Aβ_42_ peptides accumulation between AsymAD and AD cases **(Fig. 1E)**. However, a significant increase in Aβ_40_ levels among AD subjects in comparison to AsymAD and healthy control was observed **(Fig. 1F)**. Considering that Aβ_42_ aggregates are the major components of amyloid plaques in AD patients and Aβ_40_ aggregates predominantly accumulates in the blood vessels during Cerebral Amyloid Angiopathy (CAA)^41, 42^, the high levels of Aβ_40_ in AD cases compared to AsymAD cases could be due to CAA, which was mainly observed in AD (**Supplementary Fig. 2**).

**Figure 1:**
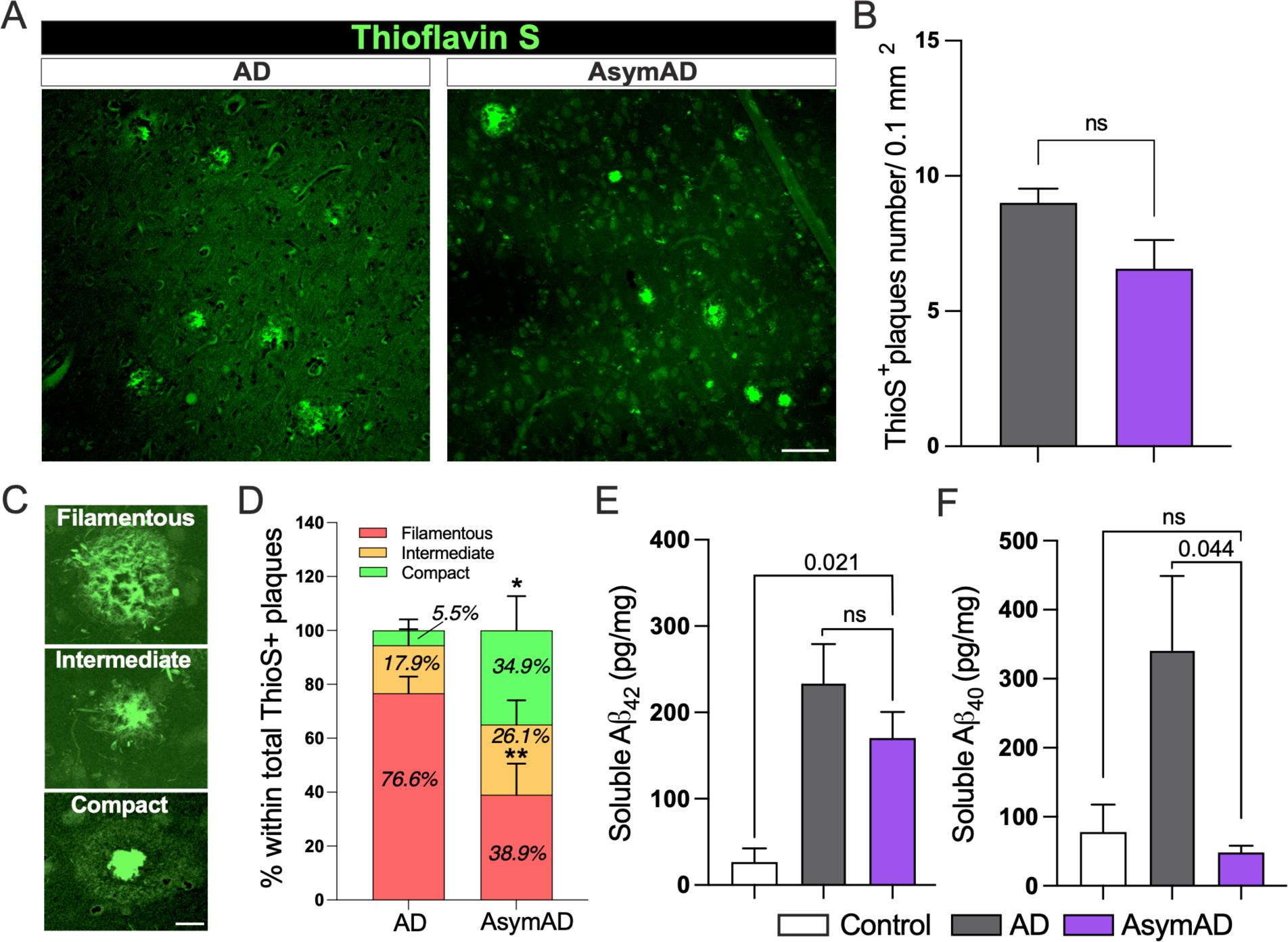
Morphologically distinct proportion of thioflavin S amyloid plaques and Aβ isoform levels in AsymAD cases compared to Alzheimer’s disease cases. **A.** Thioflavin S staining of middle frontal gyrus from AD and AsymAD subjects. Scale bar: 40 μm **B.** Quantification of total number of thioflavin S plaques. Data is shown as mean ±*SEM*, unpaired Student’s t-test, *n=14* cases per condition (n.s; 0.156) **C.** Representative morphologies of each plaque type, classified based on circularity score. Scale bar: 10 μm **D.** Proportion of filamentous, intermediate, and compact thioflavin S plaques were quantified. *n=14* cases per condition and 15-18 plaques per case were analyzed. Data is shown as ±*SEM,* 2-way ANOVA and Bonferroni’s multiple comparisons test (*p*-value *; 0.042 and **;0.001) **E-F.** Aβ_42_ (E) and Aβ_40_ (F) levels using Meso Scale Discovery assay in total soluble MFG extracts. Data is shown as mean ±*SEM,* one-way ANOVA, following Tukey’s multiple comparisons test (n.s.; 0.394 and n.s.; 0.952 for Aβ_42_ and Aβ_40_, respectively), *n=6-8* cases per condition.

### Microglia association around filamentous amyloid-plaques is increased in AsymAD cases

Microglial and astrocytic interactions with Aβ amyloid plaques have been associated with amyloid plaque development and neuritic damage. In this context, microglia appear crucial to the initial appearance and structure of plaques and, following plaque formation, they promote a chronic inflammatory state modulating neuronal gene expression changes in response to Aβ in AD pathology^43^. Furthermore, microglia limit diffuse plaques by constructing and maintaining dense compact-like plaque properties thereby blocking the progression of neuritic dystrophy^39, 44^. To investigate whether the differential proportion of amyloid plaques found in AsymAD cases compared to AD correlated with dysregulation of plaque-associated microglial and astrocytic responses, we performed a detailed analysis using MFG sections stained with thioflavin S together with anti-IBA1 and anti-GFAP, to visualize activated microglia and astrocytes associated with Aβ plaques **(Fig. 2A)**. A trend towards increased IBA1 and decreased GFAP overall coverage in AsymAD cases was observed, though not statistically significant (**Fig. 2B, C)**. However, when we measured the percentage of IBA1 and GFAP positive staining surrounding the previously three classified plaque types, we detected significantly higher levels of IBA1 around the filamentous plaques in AsymAD brains compared to AD **(Fig. 2D)**, whereas GFAP was found decreased in intermediate plaques **(Fig. 2C)**. No statistical changes were observed in other plaque phenotypes. Taken together, our observations indicate that there is a difference in the distribution of amyloid plaques between AsymAD and AD, with higher microglia abundance in the vicinity of filamentous amyloid plaques in AsymAD brains.

**Figure 2:**
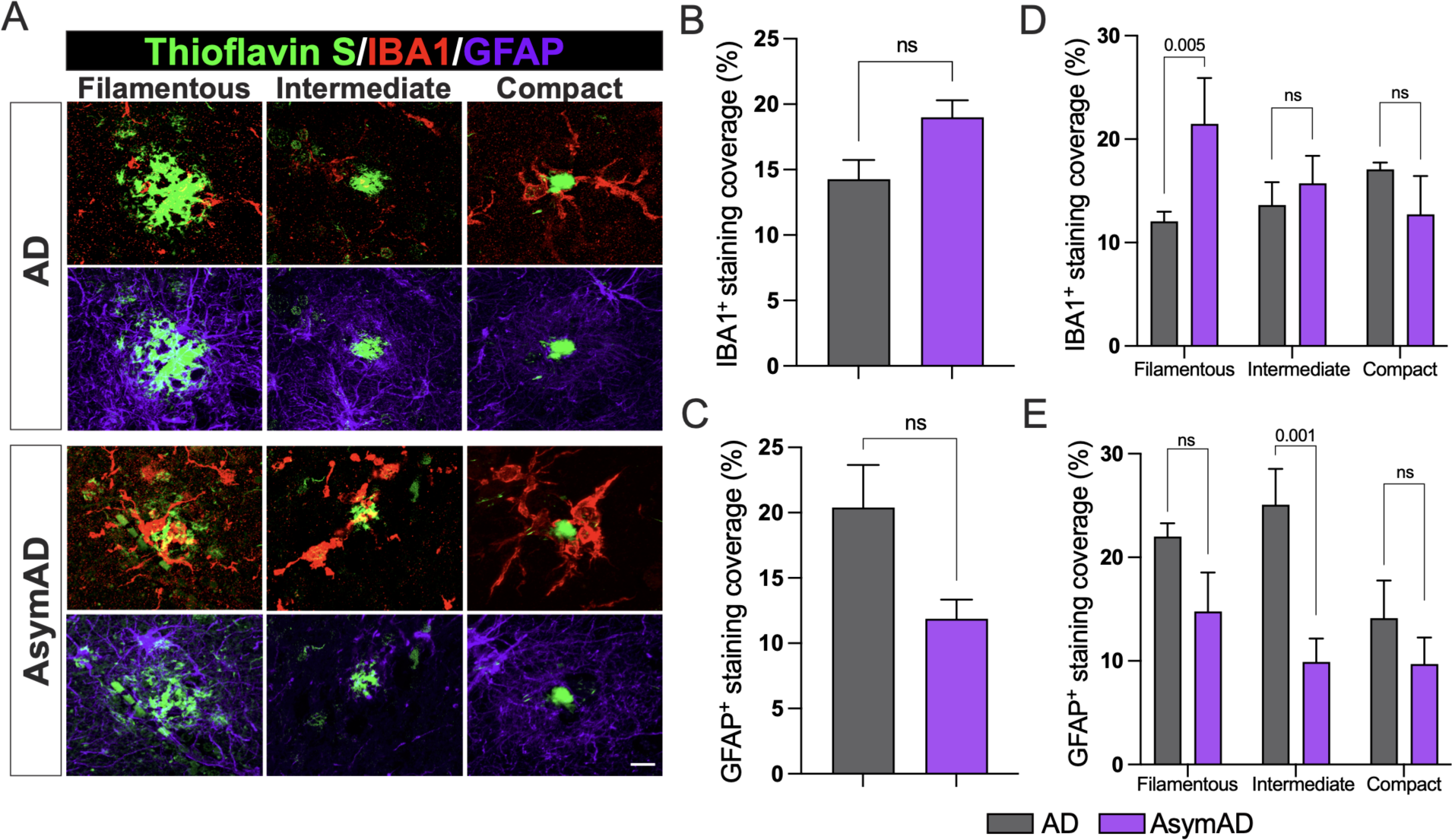
Microglia and astrocytic coverage around compact, intermediate, and filamentous amyloid plaques between AD and AsymAD cases. **A.** Staining against the activated microglia marker IBA1 (red) and the activated astrocytic marker GFAP (purple) around the three previously classified thioflavin S amyloid plaque phenotypes **B-C.** Overall IBA1 (B) and GFAP (C) coverage quantification. Data is shown as mean ±*SEM*, unpaired Student’s t-test, *n=14* cases, per condition (n.s.; 0.097 and 0.076, respectively) **D-E** IBA1 (D) and GFAP (E) coverage per plaque phenotype. Data is shown as ±*SEM,* 2-way ANOVA, following Šídák’s multiple comparisons test, *n=14* cases, per condition.

### AsymAD subjects display less pathological tau aggregation in dystrophic neurites surrounding filamentous amyloid plaques

Previous studies have demonstrated that the formation of filamentous plaques during AD pathology or aging stimulates the phosphorylation of tau within dystrophic neurites^25, 26, 45^. Moreover, amyloid-plaques create a unique molecular environment that facilitates the seeding and spread of tau pathology, leading to the formation of NFTs and neuropil threads. These highly phosphorylated tau in dystrophic neurites surrounding Aβ plaques (NP-Tau) aggregates faster and spreads more widely than tau in NFTs^24^. Microglia plays a critical role in enveloping amyloid fibrils and promoting their compaction in both AD mice models and humans, thereby avoiding axonal dystrophy and reducing tau phosphorylation in the local plaque environment^26^. As AsymAD subjects exhibited higher microglia coverage of filamentous amyloid plaques than AD subjects, we investigated whether this phenomenon could influence the formation of NP-tau in the plaque niche of AsymAD cases. We performed immunofluorescence using AT8 antibody, which recognizes the phospho sites Ser202 and Thr205 within the tau protein, and the microglial marker IBA1, in combination with thioflavin S **(Fig. 3A)**. Our findings revealed a dramatic decrease in NP-tau within the plaque microenvironment of AsymAD subjects compared to those with AD **(Fig. 3B)**. These results indicate the presence of a protective niche in the proximity of plaques in the MFG of AsymAD brains that prevents pathological tau conversion, despite the presence of toxic Aβ. Interestingly, in comparison to AD subjects, AsymAD cases did not exhibit any significant differences in intraneuronal NFTs **(Supplementary Fig. 3A-B)**. However, there was a noticeable reduction in AT8 neuritic staining **(Supplementary Fig. 3C)**, which suggests that the overall decrease in pathological neuritic tau could be attributed, in part, to the lower levels of NP-tau exhibited in the plaque microenvironment of AsymAD cases.

**Figure 3:**
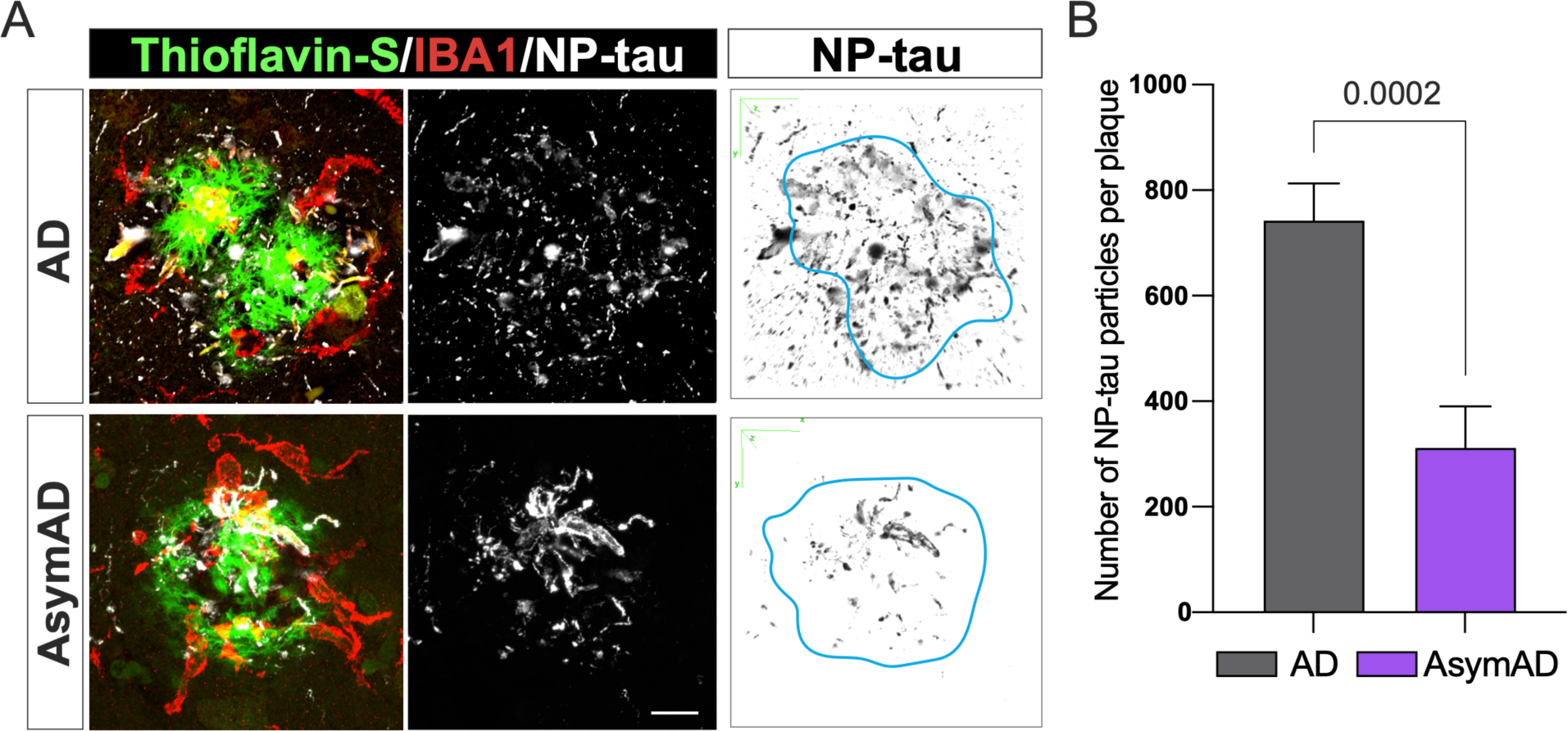
Tau aggregates in dystrophic neurites (NP-tau) surrounding filamentous amyloid plaques in AD compared to AsymAD cases. **A.** Confocal pictures of NP-Tau (AT8 – white) and microglia (IBA1 - red) together with plaques (thioflavin S - green) of AD and AsymAD cases. Black and white images are 3D reconstruction of NP-Tau particle distribution surrounding amyloid plaques. Light blue line indicates plaque location and shape in the original picture. Scale bar: 10 μm **B.** Number of NP-tau particles per plaque using the “Analyze Particles” plugin of Fiji. Data is shown as mean ±*SEM*, unpaired Student’s t-test. *n=20-30* plaques per condition.

### Seeding capacity and biochemical characterization of tau in AsymAD subjects reveals a lack of pathological tau features

Given the reduced levels of NP-tau surrounding the plaque microenvironment in AsymAD cases, our next aim was to investigate whether this phenomenon may influence soluble tau pathology. First, we evaluated the tau-seeding activity of TBS-soluble lysate from controls, AsymAD and AD cases by transfection into tau RD P301S fluorescence resonance energy transfer (FRET) biosensor cells and quantified the integrated FRET density by flow cytometry as we previously described^36, 46^. Although total tau levels between age-matched controls, AD and AsymAD cases were similar **(Fig. 4A)**, tau present in AsymAD brain lysates was not able to produce seeding activity unlike AD lysates **(Fig. 4B)**. Several groups have demonstrated that tau oligomers, which form prior to and independent of NFTs, are the toxic agents responsible to promote synaptic dysfunction in AD and drive cognitive decline^47–51^. To better understand our previous findings, we conducted a biochemical characterization of tau by immunoblot. Our results indicate that AsymAD brain lysates exhibit significantly lower levels of oligomeric tau species compared to AD cases (**Fig.4C-D**). Furthermore, tau species presented in AsymAD brain lysates are devoid of the pathological phospho-epitopes PHF1 and AT8 **(Supplementary Fig. 4A-D)**, indicating a reduction of pathological tau features in AsymAD cases. Furthermore, our laboratory has recently demonstrated that the primary source of tau seeding activity in cases of AD and progressive supranuclear palsy cases (PSP), a pure tauopathy, corresponds to soluble high molecular weight (HMW) tau complexes, making HMW tau-containing particles one of the main toxic entities^36^. To further gain insights into the size distribution of tau-seeding species in AsymAD cases, we performed size exclusion chromatography (SEC) on TBS-soluble MFG lysates. As we previously reported^36^, the tau species with the strongest seeding activity in AD cases was a HMW tau species in fraction 9 (>2,000 kDa) that represents a small percentage of total tau in the brain (**Fig. 4E and F**). Interestingly, although in AsymAD cases and controls the levels of total tau in fraction 9 are similar to AD (**Fig. 4F**), tau in AsymAD lacks seeding activity (**Fig. 4E**). These results suggest that biochemically, soluble tau in AsymAD cases resembles age-matched healthy controls rather than AD.

**Figure 4:**
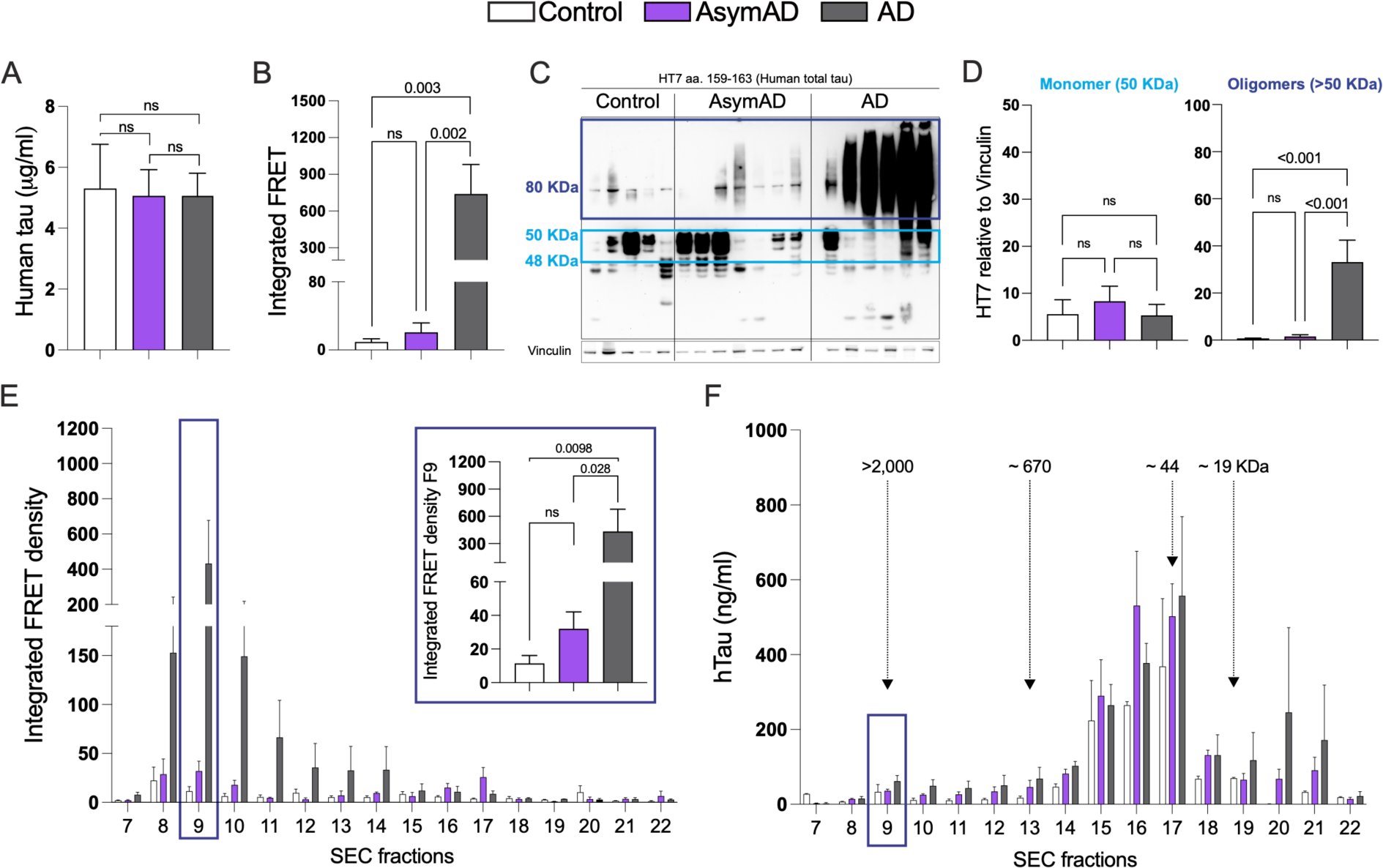
Tau seeding activity and biochemical characterization of Tau in AD, AsymAD, and age-matched control subjects. **A.** Total tau detected by ELISA in total MFG protein fractions from control, AsymAD, and AD subjects **B.** Tau seeding activity of total protein fractions **C-D.** Western blot of total HT7 and quantifications of the monomer band (between 40-50 kDa, light blue) and the oligomers bands (dark blue) (D). Data is shown as ±*SEM,* n.s. *p-*value > 0.05. Experiments were performed with *n=13-15* cases? (ELISA and tau seeding activity) and *n=5-7* cases? (Western blot) **E.** Tau seeding activity of SEC fractions. The inset shows the seeding activity of SEC fraction 9 (F9) containing the high molecular weight tau (>2000 kDa). Significance was determined by one-way ANOVA, n.s. *p-*value > 0.05 **F.** Total tau detected by ELISA in SEC fractions from MFG brain lysates. Data is shown as ±*SEM.* Experiments were performed with *n=4* cases per condition.

### Microglia from AsymAD have increased autophagy and actin-based cell motility mechanisms within the amyloid plaque microenvironment

Building upon the findings of a less detrimental plaque microenvironment, we wanted to further elucidate the underlying molecular mechanisms that may contribute to the diminished tau pathology resulting from Aβ plaques. To do so, we performed the GeoMx Hu WTA designed by NanoString, which allow us to measure over 18,000 protein-coding genes cross-referenced with the HUGO and NCBI RefSeq databases. We selected MFG sections from AsymAD and AD cases and analyzed 16-19 regions of interest (ROIs) per case. Each ROI encompassed 20-30 μm from the Aβ plaque core as well as their immediate neuronal microenvironment in the gray matter **(Fig. 5A)**. In total, we identified 696 differentially expressed genes (DEGs) by comparing ROIs of Aβ plaques in AsymAD cases (plaque-AsymAD) to the ROIs of Aβ plaques in AD cases (plaque-AD). After conducting pathway enrichment analysis on these DEGs using the GeoMx DSP analysis suite, we found that the AsymAD plaque niche was characterized by enrichment of terms related to protein translation and processing microenvironment, as evidenced by the top 20 most represented pathways **(Fig. 5B)**. Additionally, we identified enrichment for clathrin-mediated endocytosis (blue), phagocytosis-related pathways (dark cyan) and autophagy-related pathways (orange) **(Fig. 5B)**. Of the 696 DEGs up-regulated in plaque-AsymAD ROIs, 34 were in the highlighted pathways. In contrast, only 4 out of the 199 up-regulated in plaque-AD ROIs belonged to these pathways **(Fig. 5C)**. We further evaluated the expression distribution of the DEGs up-regulated in plaque-AsymAD microenvironment by plotting the normalized counts in plaque-AsymAD ROIs compared to plaque-AD ROIs. We observed a consistent and significant up-regulation of 19 DEGs in plaque-AsymAD ROIs compared to plaque-AD ROIs belonging to endocytosis, phagocytosis, and autophagy pathways **(Fig. 5D)**. Interestingly, CLU (Clusterin) and SQSTM1 (coding for p62) have been previously associated to a neuroprotective role against tau Aβ-mediated toxicity^52–55^ and reduced LAMP2 levels leads to an impaired clearance of Aβ peptides^56^. These findings are consistent with a recent study indicating that elevated expression of genes related with early stages of autophagy may be responsible for maintenance of synaptic integrity through efficient removal of tau oligomers in the hippocampus of AsymAD subjects^57^. Moreover, genetic variants associated with autophagy may play an important role in resistance to amyloid plaques and NFTs in centenarians^58^.

**Figure 5:**
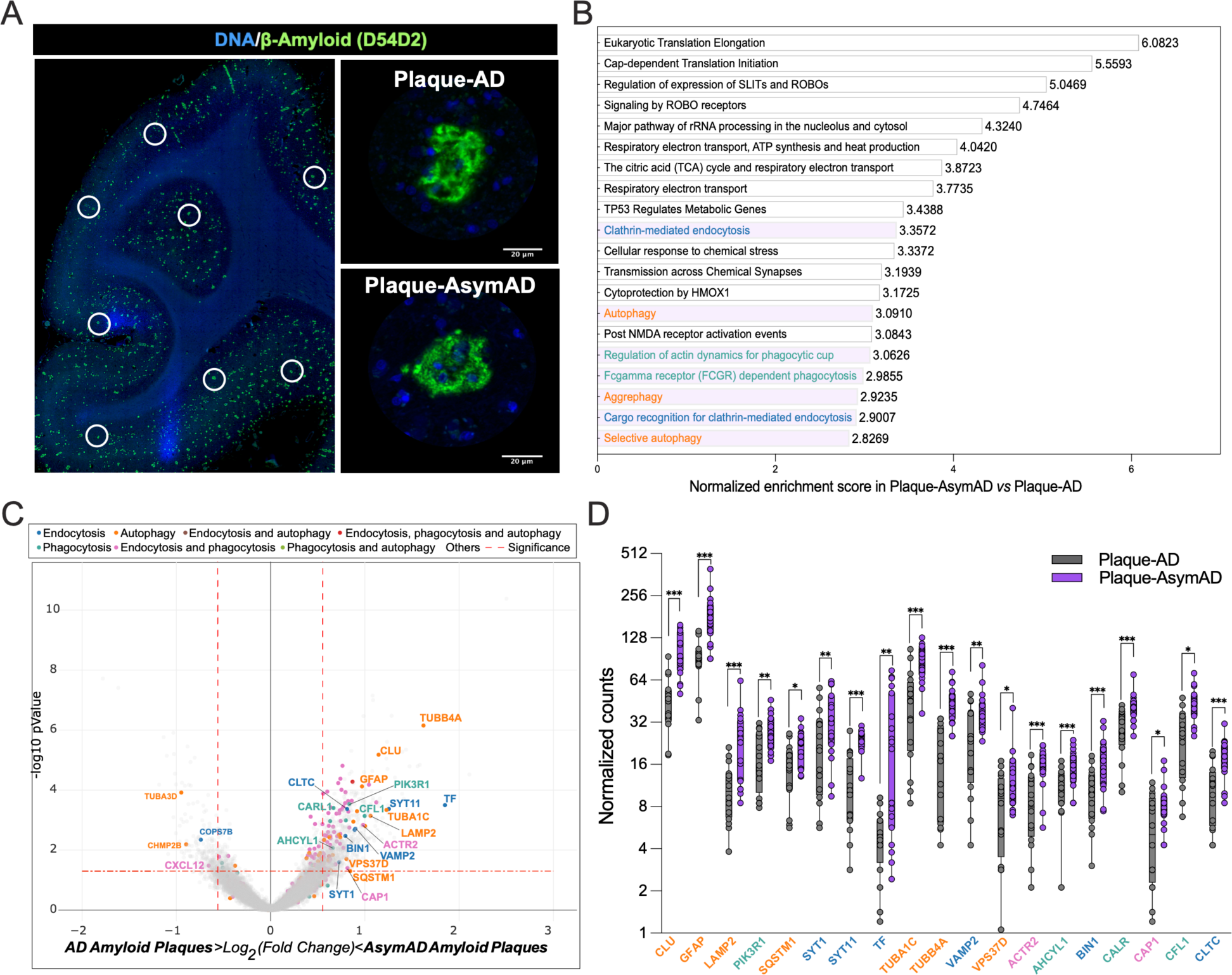
Characterization of AsymAD and AD plaque niche using spatial whole transcriptomics. **A.** After staining with Aβ-amyloid (green) and DNA (blue), areas of illumination (AOIs) containing Aβ-amyloid plaques in AD and AsymAD cases were selected. *n=2* plaques? per case. **B.** Top 20 pathway enrichment signatures in AsymAD amyloid plaques AOIs. *p*-value < 0.001 adjusted for multiple analyses using Benjamini-Hochberg procedure with a false discovery rate (FDR) of 0.01 **C.** Volcano plot of AsmAD vs AD differential expression, highlighting genes from the pathways shown in *B*. Vertical dashed lines indicates a fold change over 1.5 (log_2_ FC = *±* 0.58) and horizontal dashed indicates *p* value of 0.05 (-log_10_ *p*-value = 1.3). **D.** Normalized counts of the genes highlighted in C. Data is shown as ±*SEM,* 2-way ANOVA, following Šídák’s multiple comparisons test, one dot represents one AOI. *n=16-*18, per condition (*p*-value ***<0.0001, **;<0.0001 and *;<0.05).

Given to the crucial role of LAMP2 in the fusion of the autophagosome with the lysosome leading to cargo degradation^59^, we focused on investigating microglial LAMP2 levels and distribution within the amyloid-plaque microenvironment as well as in regions free of plaques **(Supplementary Fig. 5A-C)**. AsymAD cases presented higher levels of LAMP2 within microglial cells in the vicinity of amyloid plaques **(Supplementary Fig. 5A-B**), whereas no differences between groups were found in plaque-free areas **(Supplementary Fig. 5C-D)**. Also, we observed an accumulation of LAMP2 in the surroundings of amyloid plaques and outside microglia in AD cases. This abnormal distribution of LAMP2, indicative of dystrophic neurites^60^, was not observed in the AsymAD plaque microenvironment **(Supplementary Fig. 5A)**.

Ten DEGs were identified as being associated with endocytic (blue) and phagocytic (dark cyan) pathways. Although our data support previous evidence in which phagocytosis and endocytosis might underlie synaptic resilience in AsymAD individuals^29, 61^, the literature strongly suggests their important role in microglial motility through their interaction with actin. Specifically, ACTR2 (coding ARP2), CFL1 and CAP1, plays a significant role in actin remodeling, enabling both baseline movement (ruffling or branching) and chemotactic motility (migration)^62–64^. While ARP2 (Actin related protein 2) is part of the Arp2/3 complex, and its role is to engage with actin to start the formation of a new filament branch, CFL1 (Cofilin 1) depolymerizes filaments to make actin available for the formation of new actin structures mediated by CAP1 (Cyclase associated actin cytoskeleton regulatory protein 1)^64^. Considering that a reduction in ARP2 in excitatory synapses has been linked to AD and Down syndrome^65^ and that the Arp2/3 complex is critical for maintaining microglial morphology, branching and motility^63, 66^, we aimed to determine if ARP2 protein levels are increased in the plaque-AsymAD microenvironment, supporting the spatial transcriptomic data. First, using antibodies against ARP2 (green) and IBA1 (red) we evaluated the levels and distribution of ARP2 in the proximity of amyloid plaques (blue and labeled as P), and in microglia **(Fig. 6A)**. When we analyzed the overall fluorescence intensity of ARP2, we found a significant increase in plaque-AsymAD compared to plaque-AD **(Fig. 6B)**. Furthermore, the mean of ARP2 immunoreactivity was increased in the area occupied exclusively by IBA1 positive staining in the AsymAD plaque microenvironment, suggesting that there is an increase of ARP2 levels in microglia surrounding the amyloid plaque in AsymAD compared to AD **(Fig. 6C).** The evaluation of the colocalization between ARP2 and IBA1 using Pearson’s correlation coefficient analyses indicated that more of the 50% of the ARP2 staining is within IBA1 area, suggesting that the overall increase of ARP2 levels in AsymAD plaque microenviroment is mainly due to increased expression in microglia surrounding plaques **(Fig. 6D)**. We also analyzed ARP2 immunoreactivity in plaque-free areas **(Fig. 6E)**. Although we did not find differences in the ARP2 intensity per IBA1 cells between AsymAD and AD, AsymAD cases showed significantly higher levels of ARP2 compared to controls **(Fig. 6F)**, suggesting that AsymAD cases may possess higher baseline levels of ARP2. Overall, these data suggest that in AsymAD patients, microglia have a more efficient autophagy mechanism and a significant upregulation of actin-based motility genes that may heighten microtubules dynamics, facilitating efficient migration towards the vicinity of the plaque and promoting elongation of microglial branches to enhance its engagement with the plaque.

**Figure 6:**
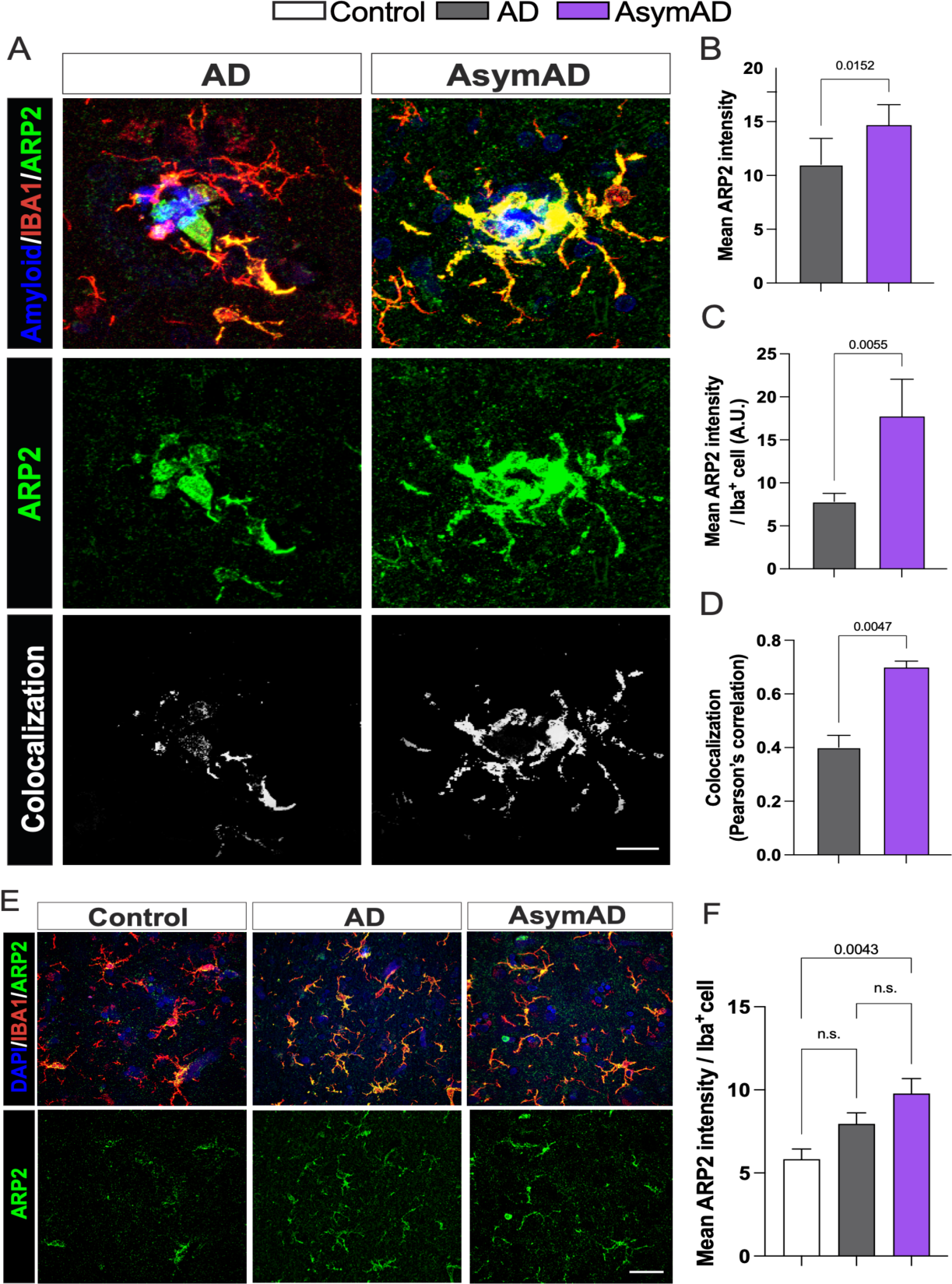
ARP2 levels are enriched in the plaque microenvironment of AsymAD cases. **A.** Staining against ARP2 (green), IBA1 (red) and DAPI, to identify amyloid-plaques and cell nuclei, respectively in control, AsymAD, and AD cases. Colocalization between ARP2 and IBA1 pixels are shown in white **B-D.** Quantification of overall mean ARP2 intensity within 50 μm of the core plaque (B) Mean ARP2 intensity per IBA1 cell (C) and Pearson correlation of ARP2 and IBA intensity (D) **E.** ARP2 and IBA1 immunostaining in areas free of amyloid-plaques **F.** Quantification of mean ARP2 intensity per IBA1 cell. Data is shown as mean *± SEM.* In A, B and C, 50-60 plaques were analyzed per condition, *n=6* cases per condition. Significance was determined by Mann-Whitney test. In F, a total of 90-170 microglia cells were analyzed per condition, *n=6* cases per condition. Data is shown as ±*SEM,* One-way ANOVA, following Tukey’s multiple comparisons test (n.s.; 0.2037 and n.s.; 0.1875).

## Discussion

Resilience has been defined as the capacity of the brain to maintain cognition and function in aging and disease based on underlying cognitive reserve, brain reserve and/or brain maintenance^67^. In this study, we have uncovered novel mechanisms that could contribute to the cognitive reserve in AsymAD individuals, associated with a distinct cell response comprised of high expression of autophagy-related genes such as LAMP2 and actin-based cell motility-related genes such as ARP2 that may stimulate the deposition of Aβ into dense-core plaques and prevent the amyloid-driven tau pathogenesis within the amyloid-plaque microenvironment of filamentous plaques.

Aβ amyloid plaques are composed of both Aβ42, being the primary component^68^, and Aβ40, which is the major constituent of amyloid deposits in the cerebral vasculature too^69^. Aβ peptides of varying lengths can transform into oligomeric and fibrillar forms, leading to the eventual formation of amyloid plaques^70^ which are classified according to their morphology^71^. The decreased filamentous/diffuse plaque type and increased compact plaque type in AsymAD cerebral cortex, raised the question whether Aβ peptide and/or the microenvironment play roles in the differences in plaque morphology distribution between AsymAD and AD cases. A possibility is that the environment in which the amyloid plaques are forming could modulate Aβ plaque shape. In this context, multiple evidence suggests that microglia around plaques is a key factor to regulate amyloid plaques dynamics and morphology in mice models of AD. Initial observations revealed that reactive microglia encircled amyloid plaques and sequester Aβ amyloid within their cytoplasm *in vitro*^72, 73^. Later studies using AD mice models supported these observations; in the CRND8 and 5xFAD models it has been shown that microglia form a tight barrier around plaques preventing their growth, and in regions lacking microglia processes, the neuritic dystrophy was more severe, these protective mechanisms were reduced with age^74^. Casali et al. showed that pharmacological depletion of microglia in ten-month-old 5xFAD mice reduced plaque burden, the remaining plaques exhibit an increase diffuse-like and fewer compact-like shapes, together with an increased in dystrophic neurites^44^. Similarly, Spangenberg et al. observed a reduction in dense-core plaques in the cortex of the same AD mice model following chronic administration of an inhibitor of microglia proliferation^43^. Using the APP/PS1 AD mice model, Huang et al. showed that the genetic ablation of tyrosine kinase TAM receptors inhibits microglia phagocytosis with a decrease in dense-core plaque in cortex and hippocampus after 12 months old, these changes were not due to any change in the production of Aβ peptides^39^. Approaches in which phagocytic microglia associated to plaques is decreased by ozone, also demonstrated exacerbation in dystrophic neurites^75^. These findings suggest that microglia may be a critical regulator of plaque conformation, and dense-core Aβ plaques do not form spontaneously but are instead constructed from loosely organized Aβ material by phagocytic microglia. Moreover, the importance of microglia-plaque association lies in their ability to restrain amyloid plaques from causing synaptic damage. The variability in microglia and astrocyte distribution based on plaque phenotype in AsymAD compared to AD cases provides insight into the complexity of amyloid-associated glial response. One study showed that reactive astrocytes and activated microglia respond differently to Aβ plaque formation: while microglia respond directly to the presence of plaques, astrocytes are associated with neuritic damage that occurs when synapses are already dysfunctional^76^. Therefore, a stronger microglia barrier surrounding plaques in the cortices of AsymAD subjects may confer a mechanism of protection against synaptic derangement and neuritic damage, which in turn could mitigate an astrocytic response.

Previous studies in AD mice models support the importance of the Aβ plaque microenvironment in promoting the pathological conversion of tau. In a mouse model of amyloidosis, it has been observed that Aβ plaques create a unique environment that triggers tau phosphorylation within dystrophic neurites^25^. Furthermore, He et al. reported that when human AD-derived tau was injected into the plaque-bearing 5xFAD mice model, Aβ plaques facilitated the conversion and seeding of pathological tau in dystrophic neurites during the early stages of the pathology. This tau propagation could potentially occur through axonal transmission to neuronal soma and dendrites, ultimately leading to the formation of NFTs^24^. However, the formation of NFTs is not entirely dependent on Aβ plaque–mediated tau pathogenesis^24^, which could explain why AsymAD cases still exhibit a considerable presence of NFTs independent of the NP-tau. Our data indicates that AsymAD cases exhibited an overall deficiency in oligomeric and soluble phospho tau species. Also, our findings provide, to our knowledge, the first evidence that soluble tau species in AsymAD individuals are not able to initiate seeding as observed in AD^36^.

The GeoMx WTA revealed several genes associated with cell engulfment (endocytosis and phagocytosis) and autophagy within the plaque AsymAD microenvironment. Abundant evidence has shown that autophagy and microglial phagocytosis are impaired in AD and aged brains^77–80^. Moreover, it has been reported that genetic variants of genes related with autophagy functions are involve in resilience against AD neuropathology ^58, 61, 81^. However, how these processes contribute to AD resilience remains an area of active investigation but has not yet been extensively explored. Previous studies in AD mice models support the idea that more efficient microglia confer protection against amyloid plaque toxicity by phagocytic activity. In Trem2 or Dap12 haplodeficient mice and humans with the R47H mutation in the TREM2 gene, microglia had a markedly reduced ability to envelop amyloid deposits, decreasing compact plaques phenotypes and increasing amyloid fibrils surface of exposure to adjacent neurites, which was found to be associated with tau hyperphosphorylation^26^. Moreover, a recent study found a high abundance of activated microglia cells in the cortices of a cohort of non-demented individuals with AD pathology (referred as NDAN). The authors also reported higher levels of the microglia phagocytic complex TREM2/DAP12 in relation to Aβ amyloid plaques in NDAN compared to AD cases and concluded that microglia surrounding Aβ plaques in NDAN subjects are hyperactive and more effective at recognizing damaged synapses with a greater phagocytic capacity than microglia in AD^29^.

Microglial function is highly dependent on baseline motility, which consists of the extension, retraction, and movement of the microglial processes^82, 83^, allowing the formation of a membrane ruffling. In APP/PS1 mice, a study using two-photon imaging showed that the ability of microglia to generate new branches and their speed was impaired after a laser insult^84^. Moreover, aging led to a reduction of microglia adhesion and migration to fibrillar Aβ in WT and APP/PS1 mice^85^. At a molecular level, a reorganization of the cytoskeleton is necessary to carry out these processes^64^; ARP2, CFL1 and CAP1 are proteins related to engulfment processes through an actin-based motility mechanism^62, 64^. It has been previously shown an overall reduction of ARP2 content in human AD parietal cortex tissue^65^. While there is not clearly evidence of CFL1 and CAP1 changes in AD^62, 86^, CFL1 is the one of the main components of the actin/cofilin rods, which are insoluble aggregates that lead to neurodysfunction^87, 88^

Interestingly, we found that in plaque-free areas, microglia from AsymAD cases exhibit significantly higher levels of ARP2 compared to healthy controls. This finding led us to speculate that AsymAD subjects may have higher basal level of ARP2 in microglia compared to AD, even before plaque development, and may possess the ability to significantly increase ARP2 expression in the presence of AD pathology helping to counteract synaptic deterioration. It is worth mentioning that ARP2 is not solely expressed in microglia, therefore we cannot rule out the effects in other cell types^89, 90^. Our GeoMx WTA also showed an enrichment of synaptic and neurotransmission release-related genes (VAMP2, SYT1, SYT11 and BIN1), as well as coding genes for microtubule proteins within the axons (TUBA1C and TUBB4A) in plaque-AsymAD-plaque compared to plaque-AD. It is well established that the absence of dementia in AsymAD individuals is partly due to synaptic preservation and hence, neuron survival^14, 91^.

Finally, this study suggests a potential mechanism by which AsymAD brains resist or slow down the pathological processes that lead to synaptic dysfunction mediated by the amyloid plaque niche. Further mechanistic experiments will be necessary to explore the underlying molecular mechanisms of the genes here described in synaptic protection and their contribution to AD resilience.

## Conclusion

Our findings reveal a novel mechanism by which AsymAD subjects can maintain normal cognition and achieve resilience against AD. We have observed that microglia cells in AsymAD brains display more efficient chemotactic motility in comparison with AD brains, being able to reach, remodel their branches and embrace the amyloid plaque, followed by a facilitated engulfment and clearance of toxic Aβ aggregates, which may mitigate the Aβ-associated tau pathogenesis decreasing tau seeding species. Our discoveries have important implications for the development of interventions to halt synaptic damage in AD and forestall subsequent cognitive impairments and dementia.

## Author Contributions

N.J-G and C.L-R designed the study and wrote the manuscript. J.R. and J.T. provided human samples and performed neuropathology diagnosis. N.J-G performed histological, biochemical, and seeding experiments. N.J.-G and Y.Y. performed size exclusion chromatography and seeding experiments of S.E.C. fractions. N.J-G and P.M. performed immunofluorescence and quantitative analyses. H.K. performed MSD experiments. J.T. contributed to interpretation of histopathology data. N.J.-G performed the whole spatial transcriptomics and analyses. T.S.J. and J.Z. contributed to spatial transcriptomics analyses. C.L.-R, J.T. and J.K. critically revised the manuscript and interpretation of the data. All authors read and approved the final manuscript.

## Acknowledgements

We would like to acknowledge all brain donors and their caregivers, and the Johns Hopkins University Morris Udall Parkinson’s Disease Center of Excellence (NINDS P50NS38377), Alzheimer’s Disease Research Center (NIH P30AG066507) and BIOCARD (NIH U19AG033655 to J.T.). This study was funded by NIH/NINDS 1R01NS119280, NIH/NIA 1R01AG059639 and AARGD-591887 grants, the Department of Defense award AZ180006 to C.A.L.-R., NIH/NIA R21AG075541 to C.A.L.-R. and J.Z., 1R01GM148970 to T.S.J and J.Z. The Tau consortium Leadership award by the Rainwater Foundation to N.J.-G. This publication was also supported by the Sara Roush Memorial Fellowship in Alzheimer’s Disease, the Indiana Alzheimer’s Disease Research Center, and the Stark Neurosciences Research Institute, and made possible by the Indiana Clinical and Translational Sciences Institute, funded in part by grant # UL1TR002529 from the National Institutes of Health, National Center for Advancing Translational Sciences. The funders had no role in study design, data collection and analysis, decision to publish or preparation of the manuscript. We thank the Biomarker core at the Stark Neurosciences Research Institute and Dr. Louise Pay for constant feedback on the manuscript.

---

**Supplementary Figure 1:**
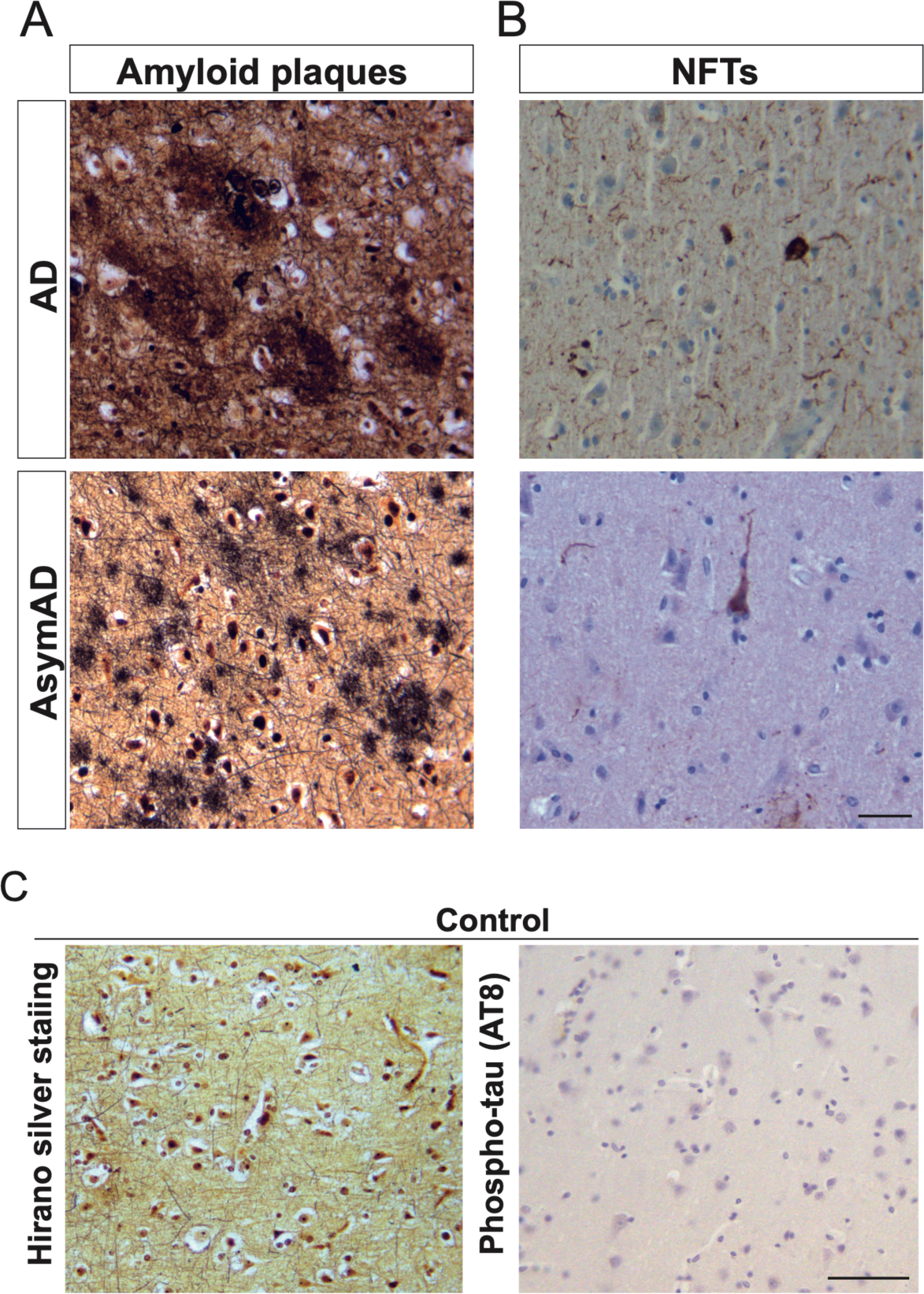
Middle frontal gyrus autopsies in non-demented subjects with high loads of Alzheimer’s disease pathology. **A-B** Histopathology performed in AD and AsymAD cases confirmed the presence of amyloid plaques by Hirano silver staining (A) and neurofibrillary tangles (NFTs) by AT8 **(B)**, an antibody which recognizes the pathological phospho-sites in Ser202/Thr305 of tau protein. Representative images showing cases with a CERAD between B-C and a Braak and Braak score of 4-6. Scale bar: 20 μm. **C.** No reactivity for none of these pathological markers was found in age-matched control subjects. Scale bar: 50 μm.

**Supplementary Figure 2:**
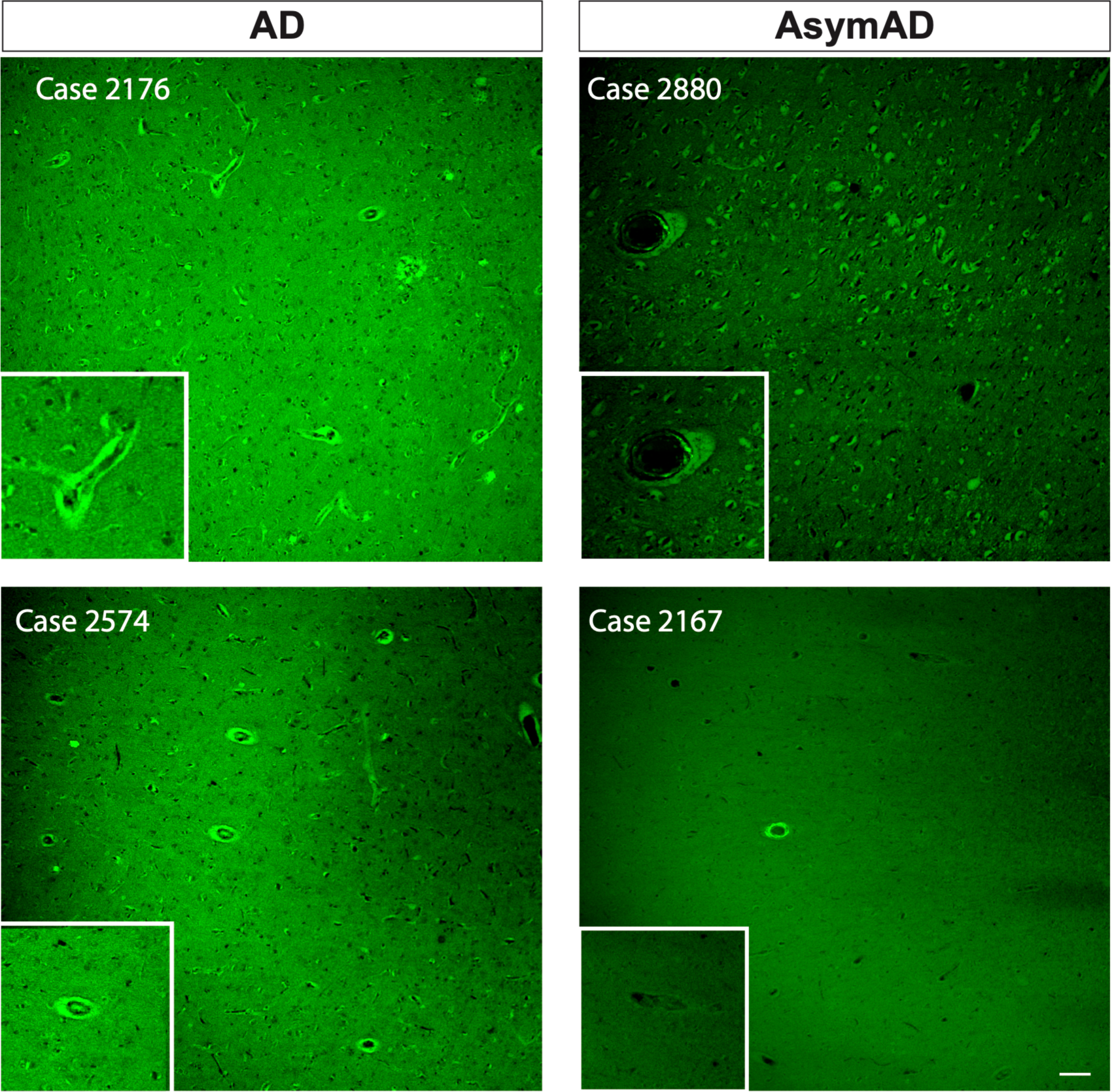
Detection of vascular amyloid aggregation using Thioflavin S in AD and AsymAD cases. The insets show structures recognizes as vasculature in the MFG of AD and AsymAD cases. Scale bar: 50 μm.

**Supplementary Figure 3:**
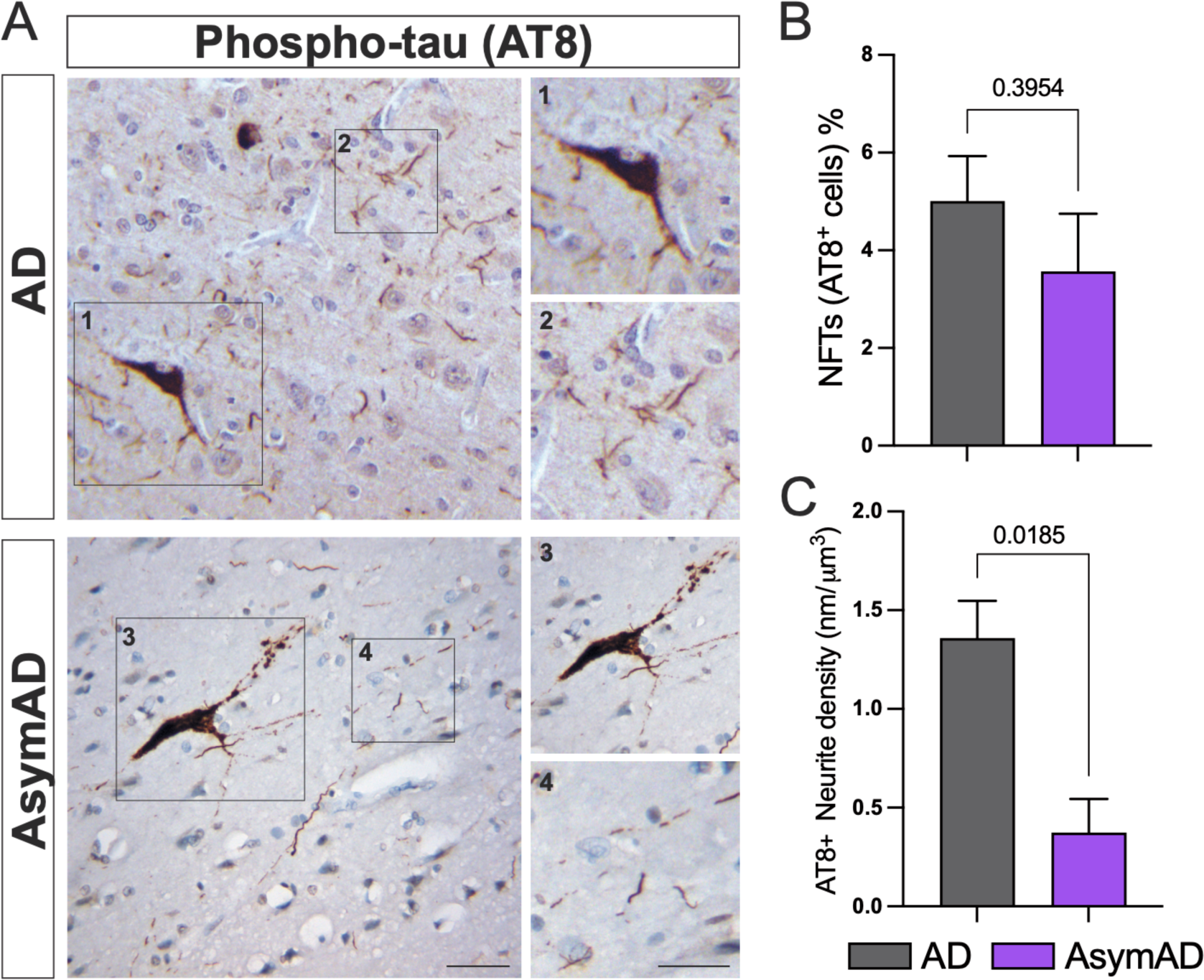
AsymAD cases show similar numbers of NFTs compared to AD, but reduced AT8 positive (AT8^+^) neuritic staining. **A.** phospho-tau staining in AD and AsymAD subjects. Both representative figures were cases classified as CERAD C and Braak and Braak score of 6. Scale bar: 20 μm and 10 μm (insets) **B.** Quantification of AT8^+^ neurites and C. percentage of NFTs/AT8^+^ cell. Data is shown as mean *± SEM,* Mann-Whitney test, *n=4-6* per condition.

**Supplementary Figure 4:**
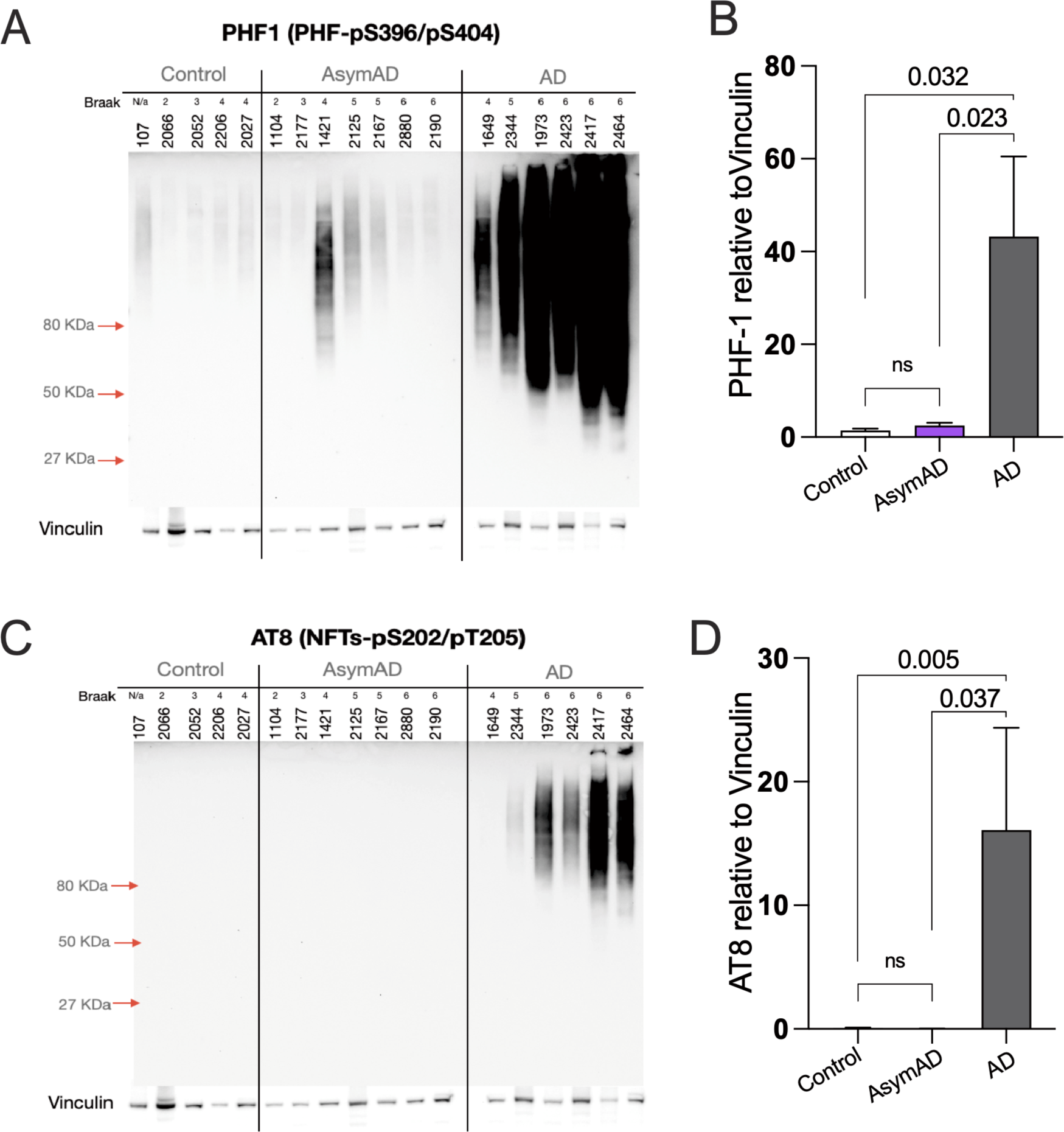
Tau species in total soluble and insoluble extracts of AD, AsymAD and aged-matched controls. **A-B.** Western blot against specific PHF1 (pTauS396/S404) (A) and AT8 (pTauS202/Thr305) (C) and its respective quantifications (B-D).

**Supplementary Figure 5:**
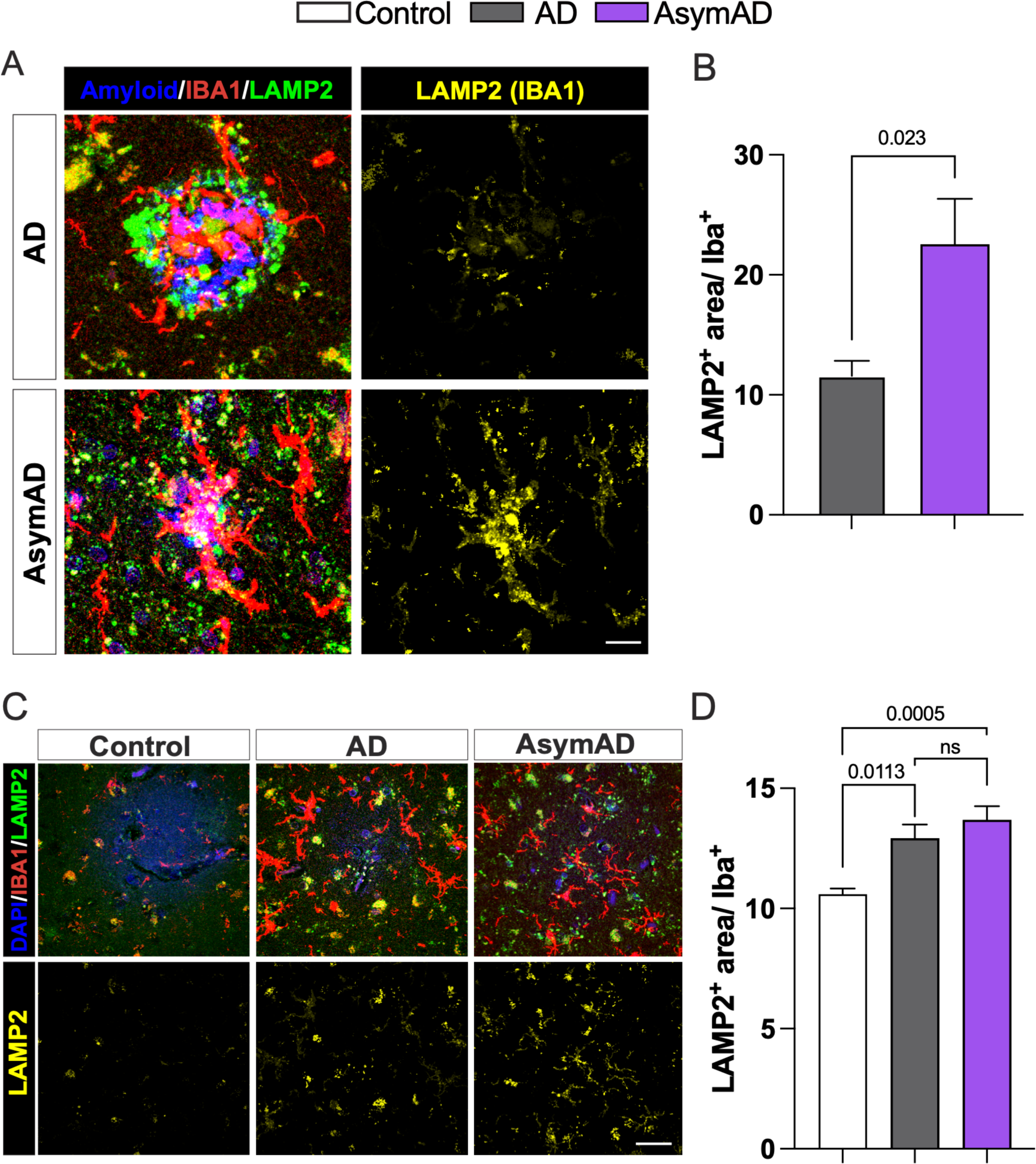
Increased LAMP2 levels in plaque-AsymAD microenvironment. **A.** Staining against LAMP2 (green), IBA1 (red) and DAPI, to identify amyloid-plaques and cell nuclei in AsymAD and AD cases **B.** Quantification of LAMP2 positive area within IBA1 area **C.** LAMP2 and IBA1 immunostaining in areas free of amyloid plaques in control, AD and AsymAD cases **D.** quantification of LAMP2 immunoreactivity in IBA1+ area. Data is shown as mean *± SEM.* In B, *n=6* cases, per condition and 22-28 plaques were analyzed. Significance was determined by Mann-Whitney test. In F, *n=6* cases, per condition were analyzed. Data is shown as ±*SEM,* One-way ANOVA and Tukey’s multiple comparisons test (n.s.; 0.5432)

